# AstroDot: a new method for studying the spatial distribution of mRNA in astrocytes

**DOI:** 10.1101/765784

**Authors:** Marc Oudart, Romain Tortuyaux, Philippe Mailly, Noémie Mazaré, Anne-Cécile Boulay, Martine Cohen-Salmon

**Affiliations:** “Physiology and Physiopathology of the Gliovascular Unit” Research Group; Center for Interdisciplinary Research in Biology (CIRB), College de France, Unité Mixte de Recherche 7241CNRS, Uniteé1050 INSERM, PSL Research University, Paris, France; Orion Imaging Facility

**Keywords:** Astrocyte, Microglia, Hippocampus, mRNA, *In situ* hybridization, Immunofluorescence, ImageJ, Gfap α, Gfap δ, Rpl4: APPswe/PS1dE9, Alzheimer’s disease, Astrocyte reactivity, Gfap, Iba1

## Abstract

Cells with a complex shape often use mRNA distribution and local translation to regulate distal functions. These mechanisms have recently been described in astrocytes, the processes of which contact and functionally modulate neighbouring synapses and blood vessels. In order to study the distribution of mRNA in astrocytes, we developed a three-dimensional histological method that combines mRNA detection via *in situ* hybridization with immunostaining of the astrocyte-specific intermediate filament glial fibrillary acidic protein (GFAP). Three-dimensional confocal images were analyzed using AstroDot, a custom Image J plug-in developed in-house for the identification and quantification of mRNAs in GFAP-immunolabelled astrocyte somata, large processes and fine processes. The custom R package AstroStat was used to analyze the AstroDot results. Taking the characterization of mRNAs encoding the astrocyte-specific GFAP α and δ isoforms in the hippocampus as a proof of concept, we showed that *Gfap* α and *Gfap* δ mRNAs mainly colocalized with GFAP in astrocyte processes. *Gfap* α mRNA was more abundant than *Gfap* δ mRNA, and was predominantly found in fine processes. Upon glial activation in the APPswe/PS1dE9 mouse model of Alzheimer’s disease, the same overall patterns were found but we noted strong variations in *Gfap* α and *Gfap* δ mRNA density and distribution as a function of the part of the hippocampus and the astrocyte’s proximity to beta-amyloid (Aβ) plaques. In astrocytes not associated with Aβ, Gfap α mRNA levels were only slightly elevated, and Gfap δ mRNA was distributed within the fine processes; these effects were more prominent in CA3 than in CA1. In contrast, levels of both mRNAs were markedly elevated in the fine processes of Aβ-associated astrocytes in both CA1 and CA3. In order to validate our new method, we confirmed that *Rpl4* mRNA (a ubiquitously expressed mRNA encoding the large subunit ribosomal protein 4) was present in large and fine processes in both astrocytes and microglia. In summary, we have developed a novel, reliable set of tools for characterizing mRNA densities and distributions in the somata and processes of astrocytes and microglia in physiological or pathological settings. Furthermore, our results suggest that intermediate filaments are crucial for distributing mRNA within astrocytes and for modulating specific *Gfap* mRNA profiles in Alzheimer’s disease.

## Introduction

Astrocytes are the most abundant glial cells in the brain. Although the astrocytes’ characteristics vary from one region of the brain to another, they all have a large number of processes that ramify into branches and then secondary branchlets. Hence, protoplasmic astrocytes are large, bushy-shaped cells with diameters of ∼40-60 µm and volumes of ∼10^4^ µm^3^. Each astrocyte covers a unique domain, and (in humans) contacts up to 2 million synapses ^1^. At the synaptic interface, perisynaptic astrocyte processes (PAPs) sense the extracellular interstitial fluid, take up neurotransmitters and ions ^2^, and release neuroactive factors ^3, 4^. Astrocytes are also in contact with blood vessels; indeed, the latter are entirely sheathed in perivascular astrocyte processes (PvAPs) ^5^. The astrocytes at this interface modulate the integrity and functions of the blood-brain barrier, immunity ^6^, cerebral blood flow ^7^, and interstitial fluid drainage ^8^. The mechanisms underlying the astrocyte’s synaptic and vascular influence are critically important, and have attracted much research interest. Indeed, dysregulation of the astrocytes’ functions and interplay with neurons and the vascular system contributes to the development and progression of most neurological diseases ^7, 9, 10^.

Recent studies of the astrocyte’s functional polarity have suggested that mRNA distribution and local translation regulates protein delivery in space and time. In a previous study, we showed that local translation is determined in PvAPs and we characterized the locally translated molecular repertoire ^11^. Local translation has also been observed in the radial glia during brain development ^12^ and in PAPs in the adult cortex ^13^. Interestingly, these studies showed that some mRNAs were specifically present in low or high levels in the astrocyte soma or processes; hence, mRNA distribution appears to obey specific rules and meet specific needs, and may help to regulate distal perivascular and perisynaptic functions. To further characterize the mRNA distribution in astrocytes, we developed a new three-dimensional *in situ* method for identifying astrocyte mRNAs colocalized on GFAP-immunolabelled processes and quantifying these levels with dedicated bioinformatics tools. More precisely, we studied the distribution of mRNAs encoding the astrocyte-specific GFAP α and δ isoforms (generated by alternative splicing) in the CA1 and CA3 regions of the hippocampus in wild-type (WT) mice and in the APPswe/PS1dE9 mouse model of Alzheimer’s disease (AD). We further showed that our approach can be applied to microglia (immunolabelled for ionized calcium binding adaptor molecule 1 (Iba1)) and to all types of mRNA.

## Results

### *Gfap* α and *Gfap* δ mRNAs are distributed in PAPs

*Gfap* mRNAs have already been detected in distal perivascular ^11^ and perisynaptic processes ^13^ of astrocytes - suggesting that local GFAP translation regulates distal intermediate filament assembly. Although previous research focused on the canonical isoform GFAP α, at least 10 GFAP isoforms (generated by alterative mRNA splicing and polyadenylation signal selection) have been described ^14-17^. GFAP δ has received special interest because it is a tumour marker ^18-20^ and is associated with neurogenic niches ^21^. In fact, GFAP δ is encoded by the same first 7 exons as GFAP α but has a different C-terminal (Fig. 1A), allowing it to form a heterodimer with GFAP α and thus promote intermediate filament aggregation ^16, 22^. Here, we first looked for GFAP α and GFAP δ mRNAs in PAPs. Polysomal mRNAs were extracted by translating ribosome affinity purification (TRAP) from adult Aldh1-RPL10a mice, which express the ribosomal protein Rpl10a specifically in astrocytes ^23^. Extractions were performed either from hippocampus (for whole-astrocyte polysomal mRNAs) or synaptogliosome preparations (consisting of apposed pre- and post-synaptic membranes and astrocyte PAPs), in order to extract polysomal mRNAs contained in PAPs ^13, 24^. Quantitative qPCR amplification of *Gfap* α and δ mRNA was performed using specific primers (Fig. 1B). Both isoforms were detected in whole astrocytes (mean ± standard error of the mean level: 8.28±1.99 arbitrary units for Gfap α and 1.38±0.20 for Gfap δ) and in the perisynaptic processes (17.04±9.09 for Gfap α and 1.16±0.76 for Gfap δ). For polysomal mRNAs, the Gfap α/Gfap δ ratio was significantly higher in PAPs (40.09±24.27; N=3; *p*-value=0.05) than in whole astrocytes (5.81±0.67) - suggesting the predominance of Gfap α in PAPs (Fig. 1B).

**Fig. 1:**
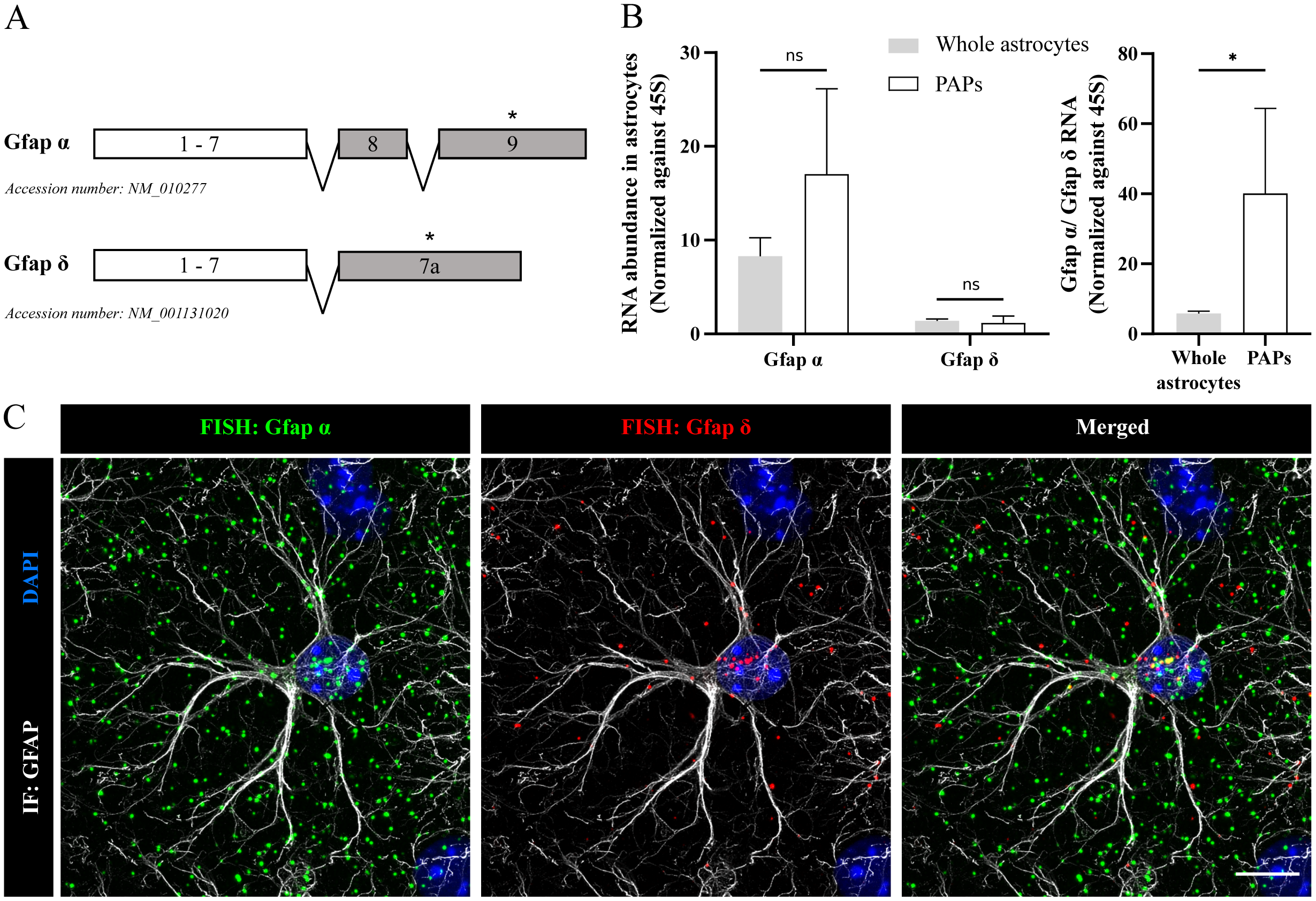
Detection of Gfap α and δ mRNAs in hippocampal astrocytes. **A**. Schematic representation of mouse GFAP α and GFAP δ isoforms. The positions of the qPCR and FISH probes are indicated by an asterisk. **B**. Polysomal *Gfap* α and δ RNA levels in hippocampal astrocytes and perisynaptic astrocyte processes (PAPs), determined by qPCR and normalized against 45S RNA. Statistical significance was determined in a one-way unpaired Mann-Whitney test; *, p<0.05; ns: not significant. **C**. Merged and separated images of a deconvoluted confocal z-stack of a CA1 astrocyte, with FISH detection of *Gfap* α (in green) and *Gfap* δ (in red) mRNAs and co-immunofluorescent detection (IF) of GFAP (in grey). The nucleus was stained with DAPI (in blue). Note the abundance of *Gfap* α mRNA FISH dots (relative to *Gfap* δ) in distal areas of the astrocyte. Scale bar, 10 μm.

We next sought to visualize *Gfap* α and δ mRNAs in hippocampal astrocytes. Fluorescent *in situ* hybridization (FISH) was performed on 30 μm thick free-floating adult mouse brain sections, using specific fluorophore-coupled RNAscope® probes against Gfap α exon 9 and Gfap δ exon 7a (Fig. 1C). Next, the astrocytes’ somata and processes were labelled by GFAP immunostaining (Fig. 1C). Importantly, the co-immunofluorescent detection of proteins depends on the preservation of their epitopes during the protease digestion step preceding FISH. We observed dense, continuous, GFAP-labelled arborizations, which indicated that our protocol preserved the GFAP epitopes. In line with the qPCR results presented above, *Gfap* α and δ mRNA FISH signals were detected as discrete dots in the soma and in distal astrocyte areas; *Gfap* α mRNA (Fig. 1B) was more abundant than *Gfap* δ, which was mainly present in the somata (Fig. 1C).

### AstroDot and AstroStat: bioinformatics tools for analyzing the mRNA distribution in astrocytes

In order to analyze the distribution of *Gfap* α and d mRNAs in astrocytes, we developed AstroDot - a dedicated ImageJ plug-in. We had two main objectives: (i) to detect mRNA FISH dots that colocalized with GFAP-immunostained astrocyte processes; and (ii) to quantify these dots and analyze their distribution in the astrocytes. Confocal images of the CA1 and CA3 regions of the hippocampus were acquired and then deconvoluted, so as to eliminate the inherent fluorescence blurring (the point spread function (PSF)) and noise, and to increase the resolution (Fig. 2A). For each individual astrocyte, regions of interest (ROIs, i.e. the soma and processes) were selected manually by assessing the GFAP (intermediate filaments) and DAPI (nuclei) staining and by defining the stack of confocal planes (Fig. 2B). AstroDot opens with a dialogue box that enables the operator to attribute a specific purpose for each of the three fluorescence channels, e.g. “DAPI” for nuclear staining, “IF” for GFAP immunofluorescence, and “Dots” for FISH dot thresholding and detection (Fig. 2C). This dialogue box also contains a “Specific mRNA” option, which can be selected when the mRNAs are expressed only in the cell type of interest. In such a case, all mRNA FISH dots are considered to belong to this cell type. The first step in the analysis was calculation of the mean GFAP immunofluorescence background, i.e. the value above which the signal was considered to be positive. In the second step, each astrocyte nucleus was defined; given that astrocytes interact with other brain cell types, some ROIs can contain more than one nucleus. AstroDot was designed to optimize the recognition of astrocyte nuclei on the basis on the GFAP immunostaining. The putative astrocyte nucleus appears in green, and any other nuclei appears in red. A dialogue box allows the operator to confirm or modify AstroDot’s automatic selection by clicking on the correct nucleus (Fig. 2D). AstroDot then starts to detect astrocyte mRNAs, based on their specific colocalization with GFAP. A distance map was used to calculate the diameter of each GFAP-immunolabelled process. Processes with a diameter greater than the minimum distance between two confocal planes (0.3 μm, in the present case), were defined as “large”, and those with a smaller diameter were defined as “fine” (Fig. 2E). The somatic domain of each astrocyte corresponded to the DAPI staining and the surrounding 2 μm space. A TIF image was generated for each ROI (Fig. 2F). The mRNA FISH dots appeared in red if they were outside astrocytes, in green if they colocalized with astrocyte large processes and somata, or in yellow if they colocalized with astrocyte fine processes (Fig. 2F). All the results were automatically entered on a table with the following items for each ROI: Image name; ROI name; Background intensity; Astrocyte volume; Dot density in astrocytes; Percentage of dots not in astrocytes; Percentage of dots in astrocyte somata; Percentage of dots in astrocyte fine processes; Percentage of dots in astrocyte large processes; Mean astrocyte process diameter. To facilitate the statistical analysis of AstroDot data, we developed an optional R package named AstroStat; it automatically calculates and compares the mean ± standard deviation values, and produces a summary report on the results.

**Fig. 2:**
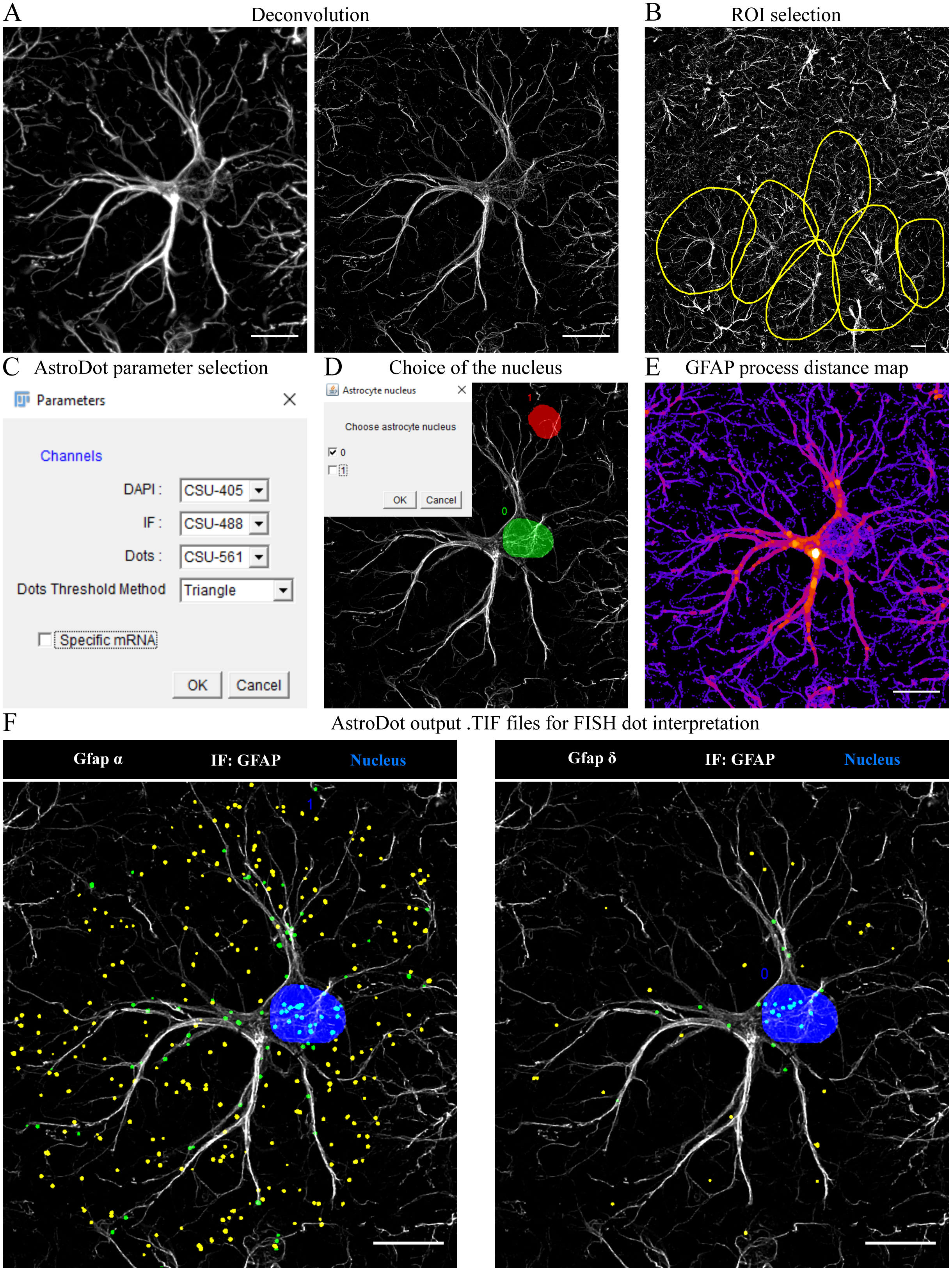
AstroDot image processing. All images correspond to a single confocal z-stack for a CA1 astrocyte. **A**. Effect of deconvolution on GFAP immunofluorescence: left panel: the raw confocal image; right panel: the deconvoluted image. **B**. Selection of ROIs (yellow circles). **C**. AstroDot dialogue box for the definition of fluorescence channels, the threshold method for FISH dots, and the choice of the “Specific mRNA” option. **D**. Detection of the astrocyte nucleus (in green) and other nuclei (in red). **E**. A heat map of GFAP immunofluorescence, used to calculate the process diameter. **F** AstroDot’s interpretation of the results for *Gfap* α and *Gfap* δ mRNAs, with the “Specific mRNA” option active. Green dots are located in the soma or large GFAP-labelled processes. Yellow dots are located in fine processes. Scale bar: 10 μm.

### Characterization of *Gfap* α and δ mRNAs in CA1 and CA3 hippocampal astrocytes from WT mice and the APPswe/PS1dE9 mouse model of AD

We analyzed the density and distribution of *Gfap* α and *Gfap* δ mRNAs in CA1 and CA3 hippocampal adult astrocytes in WT adult mice by using AstroDot’s “Specific mRNA” option (Fig. 3 and Table S2). Comparison of the astrocytes in CA1 vs. CA3 indicated that CA1 astrocytes had a slightly greater overall volume but displayed processes with the same mean diameter (Fig. 3A). The *Gfap* α/*Gfap* δ mRNA ratio was the same in the two regions (Fig. 3B). Overall, and in line with our initial qPCR analysis (Fig. 1B), Gfap α was 5.2 times more abundant than *Gfap* δ in both CA1 and CA3 (Fig. 3C). Both mRNAs were more abundant in the processes (*Gfap* α 88.5 % ± 6.7 SD in CA1 and 86.7 ± 8.1 SD in CA3; *Gfap* δ 73.4% ± 11.3 SD in CA1 and 71.5 % ± 16.4 SD in CA3) than in the soma. *Gfap* δ was more abundant than *Gfap* α in the soma and in large processes (Fig. 3D). We analyzed the data without taking account of the astrocytic-specific nature of GFAP expression (Fig. 3E). In this case, both *Gfap* α and *Gfap* δ mRNAs were colocalized on GFAP-labelled intermediate filaments in CA1 (mean ± SD: 59.5 ± 9.0 for *Gfap* α and 74.4 ± 11.0) and CA3 (62.2 ± 10.1 for *Gfap* α and 74.7 ± 12.4 for *Gfap* δ) - suggesting that the majority of *Gfap* RNAs are associated with intermediate filaments (Fig. 3E).

**Fig. 3:**
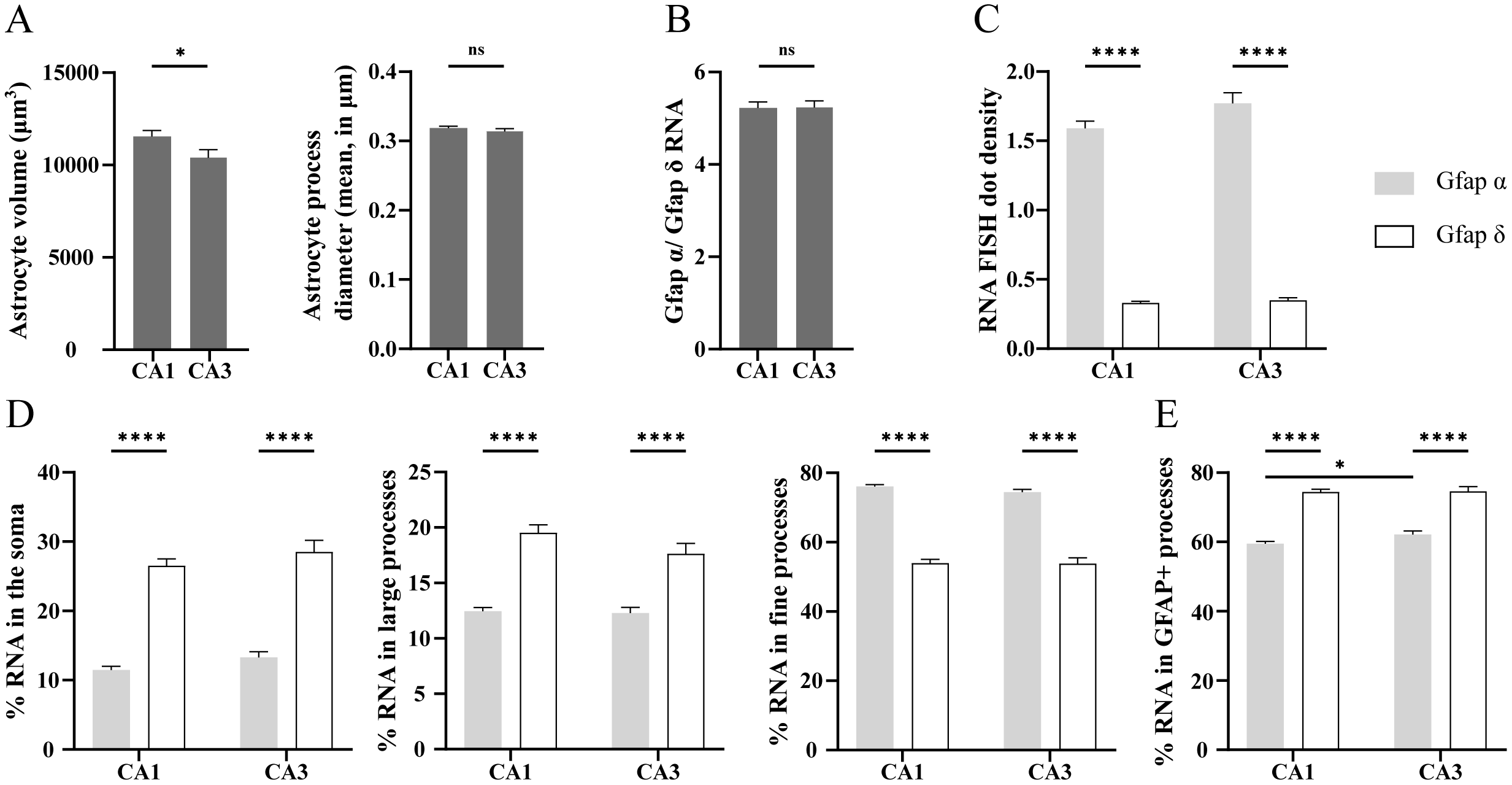
The distribution of Gfap α and δ mRNAs in CA1 and CA3 hippocampal astrocytes. **A**. Astrocyte volume, and process diameter. **B**. The *Gfap* α/*Gfap* δ mRNA ratio. **C**. Total mRNA density: number of RNA FISH dots/μm^3^ × 100. **D**. Percentages of *Gfap* α and δ mRNAs in astrocyte somata, fine processes and large processes. **E**. Percentages *Gfap* α and δ mRNA dots colocalized with GFAP when the “Specific mRNA” option was not applied. In total, 175 CA1 and 94 CA3 astrocytes were analyzed. Statistical significance was determined in two-way unpaired Student’s t-tests. *, p<0.05; ****, p<0.0001; ns: not significant.

Astrocytes are involved in neuroinflammation, and become reactive in virtually all pathological situations in the brain. Astrocyte reactivity is characterized by GFAP overexpression and subsequent morphological changes, such as process hypertrophy and remodelling ^17^. Hence, we next sought to study *Gfap* α and δ mRNAs in reactive astrocytes. We choose the example of AD, in which astrocytes undergo drastic morphological and molecular changes that perturb their physiology ^25, 26^. Using the method described above, *Gfap* α and δ mRNA FISH dots were detected on GFAP-immunolabelled sections of CA1 and CA3 hippocampus from 9-month-old APPswe/PS1dE9 mice (Fig. 4). We quantified astrocytes associated with a beta-amyloid plaque (Aβ, labelled with DAPI) or more than 30 µm from an Aβ plaque (Fig. 5 and Table S2). As reported in the literature, CA1 and CA3 astrocytes from APP/PS1dE9 mice were larger than those from WT mice (Fig. 5A) but had a slightly smaller process diameter (Fig. 5A). In astrocytes not associated with Aβ, the *Gfap* α/δ ratio was slightly but significantly higher (by a factor of 1.3) in CA1 and CA3 (Fig. 5B), with a higher *Gfap* α mRNA level in fine processes only (Fig. 4A, 5F). In contrast, the *Gfap* δ mRNA density was the same as in WT mice in CA1 and CA3 (Fig. 5C). However, the distribution of this mRNA within the astrocytes differed; levels in large processes were lower (relative to the WT) in CA1 and CA3 (Fig. 5E), and levels in fine processes were higher in CA3 only (Fig. 5F). The relative differences in mRNA levels were greater in Aβ-associated astrocytes (Fig. 4B); the density of *Gfap* α mRNAs was significantly higher than in WT cells (5.0-fold for CA1, and 4.7-fold in CA3) or in astrocytes not associated with Aβ (3.8-fold for CA1, and 3.7-fold in CA3) (Fig. 5C). The distribution of *Gfap* α mRNA also differed, with lower levels (relative to the WT) in the soma (Fig. 5D) and in large processes (only in CA1) (Fig. 5E) and higher levels in fine processes in CA1 and CA3 (Fig. 5F). The same effect was observed for the *Gfap* δ mRNA, with a greater abundance in Aβ-associated astrocytes than in WT samples (3.6-fold for CA1, and 3.5-fold in CA3) or in astrocytes far from plaques (3.3-fold for CA1, and 3.4-fold in CA3) (Fig. 5C). The redistribution was most prominent in fine processes in CA1 and CA3 (Fig. 5F). These results show that the density and distribution of *Gfap* α and *Gfap* δ mRNAs vary markedly as a function of the astrocyte’s reactivity status, the brain area, and the proximity of Aβ deposits.

**Fig. 4:**
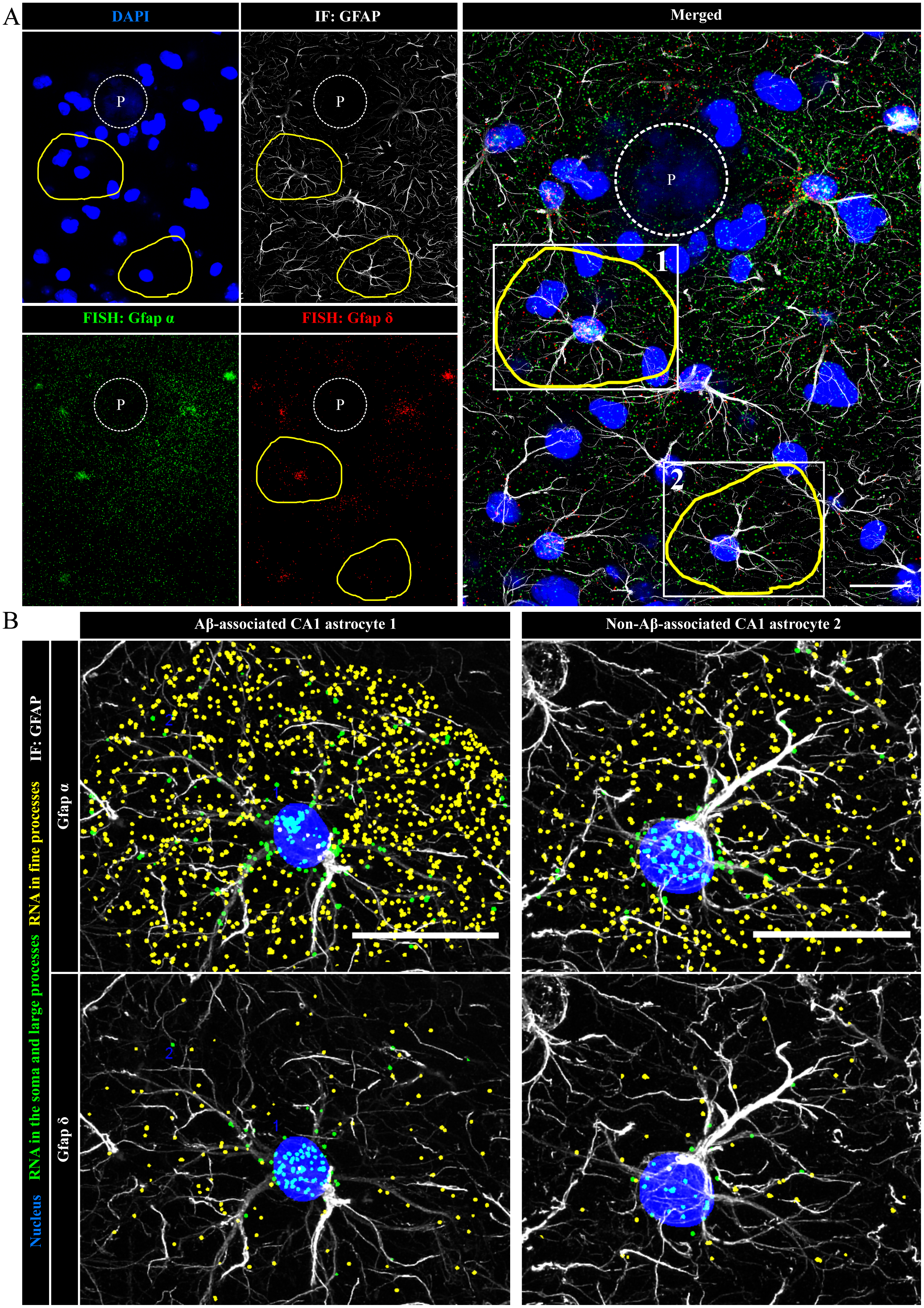
Detection of Gfap α and δ mRNAs in CA1 hippocampal APPswe/PS1dE9 astrocytes. **A**. Merged and separated images of a deconvoluted confocal z-stack of APPswe/PS1dE9 CA1 astrocytes, with FISH detection of *Gfap* α mRNA (in green) and *Gfap* δ mRNA (in red) and co-immunofluorescent detection of GFAP (in grey). The nucleus and an amyloid deposit (circled by a dotted line, and indicated by “P”) are stained with DAPI (in blue). The ROI #1 (yellow circle) is an astrocyte close to an Aβ deposit. The ROI #2 is located more than 60 μm from an Aβ deposit. **B**. TIF images of ROIs #1 and #2 for *Gfap* α and δ mRNA, as analyzed with AstroDot using the “Specific mRNA” option. Green dots belong to the soma and large GFAP-labelled immunofluorescent processes. Yellow dots belong to fine processes. Scale bar: 20 μm.

**Fig. 5:**
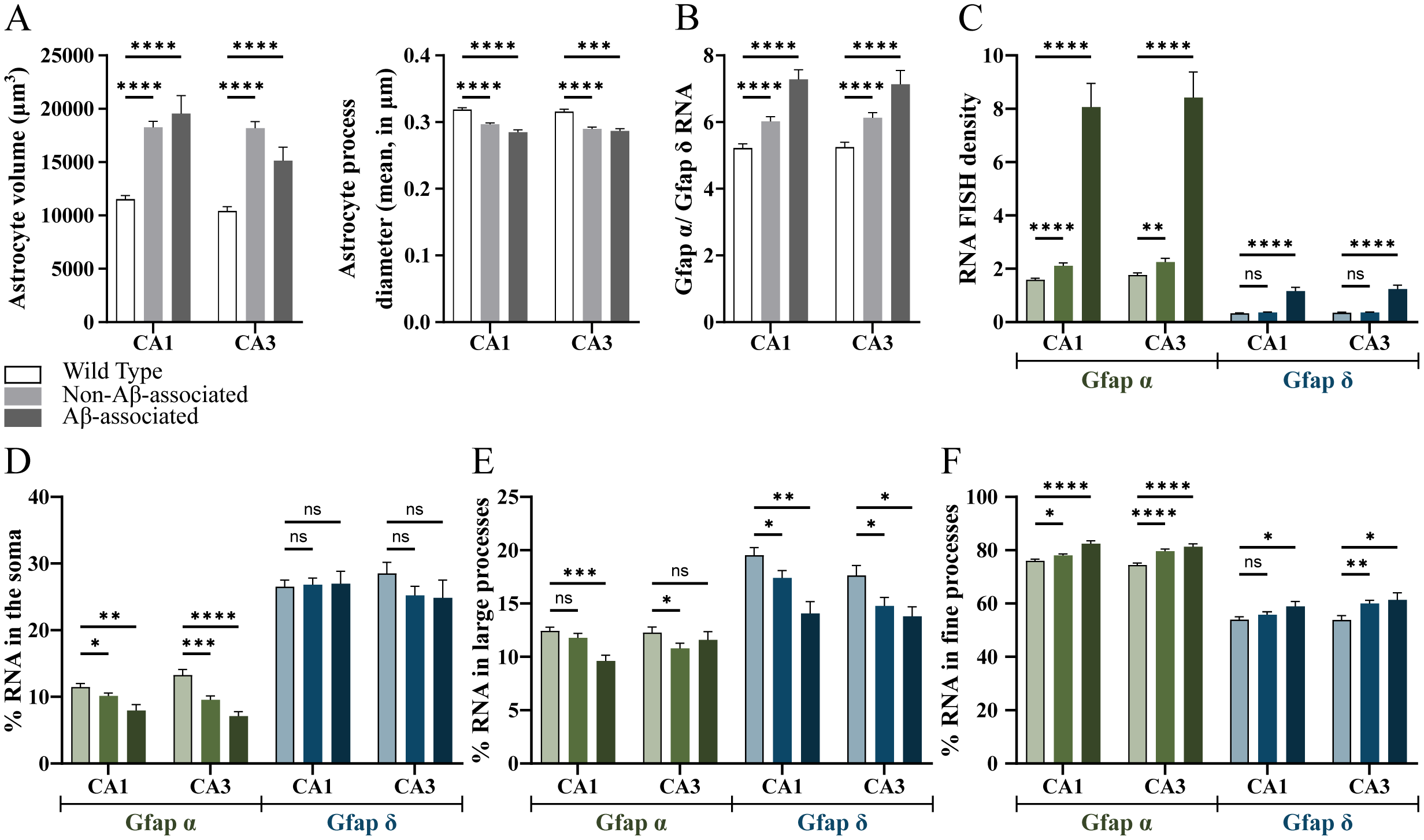
Comparison of Gfap α and δ mRNA densities and distributions in CA1 and CA3 hippocampal astrocytes from WT and APPswe/PS1dE9 mice. **A**. Astrocyte volume and process diameter. **B**. The Gfap α/Gfap δ mRNA ratio. **C**. mRNA density: number of RNA FISH dots/μm^3^ × 100. **D**. Percentages of Gfap α and δ mRNA dots in astrocyte somata, fine processes, and large processes. **E**. Percentages of Gfap α and δ mRNA dots colocalized with GFAP. Analyses were performed on 175 CA1 WT astrocytes, 94 CA3 WT astrocytes, 127 APPswe/PS1dE9 CA1 astrocytes not associated with plaques, 78 APPswe/PS1dE9 CA3 astrocytes not associated with plaques, 27 plaque-associated CA1 APPswe/PS1dE9 astrocytes, and 28 plaque-associated CA3 APPswe/PS1dE9 astrocytes. Statistical significance was determined using two-way unpaired Student’s t-tests. * p<0.05; ** p<0.001; *** p<0.001: **** p<0.0001; ns: not significant.

### Application of AstroDot and AstroStat to the analysis of ubiquitous mRNAs in astrocytes and microglia

To further validate our approach, we studied the distribution of *Rpl4* mRNA (a ubiquitously expressed mRNA encoding the large subunit ribosomal protein 4) in CA1 (Fig. 6A). Interestingly, 62.52% ± 11.77 of the *Rpl4* mRNA FISH dots were localized in astrocytes (n=67). Of these, 83.33% ± 5.41 were present in fine GFAP-immunolabelled processes, with 9.59% ± 3.45 in large GFAP-immunolabelled processes, and 7.09% ± 4.14 in somata. This result was unexpected because Rpl4 integrates into the 60S ribosome subunit in the nucleus ^27^ but was corroborated by a qPCR analysis (performed as described above) of polysomal mRNAs extracted by TRAP from adult Aldh1L1-RPL10a mouse hippocampus or PAPs; in the latter, Rpl4 was enriched 120-fold (p=0.05, n=3) (Fig. 6B). To study the distribution of non-astrocyte *Rpl4* mRNA FISH dots, we performed an additional, independent experiment by immunostaining the specific microglial marker Iba1. As with the GFAP immunofluorescence experiments, our FISH protocol was designed to protect Iba1’s epitopes; the protein was detected throughout the somata and processes (Fig. 6C). Our analysis of the distribution of *Rpl4* mRNA in microglia (n=28) indicated that 16.07% ± 4.47 of the *Rpl4* mRNA FISH dots were localized in microglial processes. Of these, 37.72% ± 9.24 were localized in fine processes, with 27.06% ± 10.78 in large processes and 35.22% ± 10.13 in somata.

**Fig. 6:**
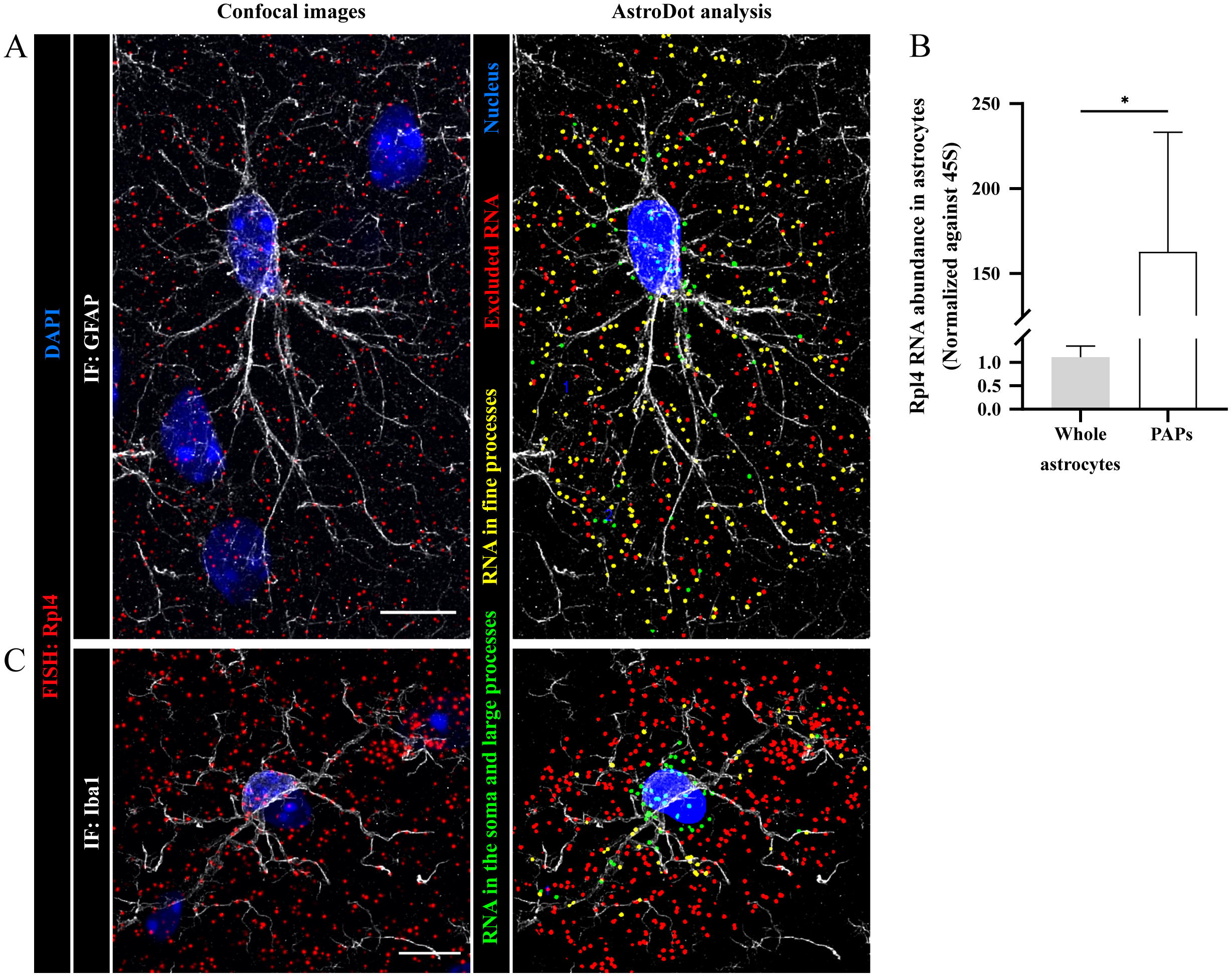
Detection and characterization of Rpl4 mRNA distribution in CA1 hippocampal astrocytes and microglia. **A**. Left: Confocal z-stack of a CA1 astrocyte with FISH detection of Rpl4 mRNA (in red) and co-immunofluorescent GFAP detection (in grey). The nucleus is stained with DAPI (in blue). Right: AstroDot analysis: Green dots are located in the soma or in GFAP-immunolabelled large processes; yellow dots are located in GFAP-immunolabelled fine processes; red dots are not colocalized with GFAP (i.e. excluded RNAs). **B**. The polysomal Rpl4 RNA level in hippocampal astrocytes and PAPs, determined by qPCR and normalized against 45S RNA. Statistical significance was determined in a one-way unpaired Mann Whitney’s test; *, p<0.05. **C**. Left: Confocal z-stack of a CA1 microglial cell with FISH detection of Rpl4 mRNA (in red) and co-immunofluorescent Iba1 (in grey). The nucleus was stained with DAPI (in blue). Right: the AstroDot analysis: Green dots are located in the soma or Iba1-immunolabelled large processes; yellow dots are located in Iba1-immunolabelled fine processes; red dots did not colocalize with Iba1 (i.e. excluded RNAs). Scale bar; 10 μm.

## Discussion

Although local translation has been recently described in astrocyte processes, tools for studying the distribution of astrocyte mRNAs were not previously available. Accordingly, we developed a co-labelling method that combined mRNA *in situ* hybridization, the immunofluorescence detection of GFAP-containing intermediate filaments on brain slices, confocal imaging, and a bioinformatics analysis of mRNA density and distribution in astrocytes.

A key technical obstacle to the implementation of this approach was the risk of protein epitope degradation during the protease digestion step that precedes *in situ* RNA hybridization. Our previous tests on transgenic hGfap-eGFP mouse brain sections (in which eGFP fills the astrocyte cytoplasm) ^28^ indicated that these adaptations were not sufficient to preserve eGFP (data not shown), and thus precluded the use of this reporter mouse strain to detect astrocytes in parallel with *in situ* hybridization. In contrast to previous reports^11, 12^, however, our protocol preserved GFAP and enabled us to perform parallel *in situ* hybridization and GFAP immunodetection. Interestingly, these conditions also allowed us to immunodetect the microglia-specific protein Iba1. It is noteworthy that GFAP is not expressed uniformly in the brain, and so GFAP immunolabelling is somewhat limited by its lack of applicability to all brain regions. Nevertheless, our optimization of the GFAP immunolabeling makes it possible to distinguish between strongly labelled astrocyte processes and their secondary extensions in regions where GFAP is highly expressed (e.g. the hippocampus, olfactory bulbs, cerebellum, and hypothalamus). Another advantage of immunolabelling GFAP and Iba1 relates to the fact that both proteins are standard markers of glial reactivity - a process initiated in response to immune attack, chronic neurodegenerative disease, or acute trauma ^29^. Hence, GFAP and Iba1 immunolabelling could therefore be used to address possible changes in mRNA distribution in reactive astrocytes and microglia, as demonstrated here in APPswe/PS1dE9 mice.

It was previously determined that GFAP immunolabeling delineates only 15% of the total astrocyte volume ^30^. Nevertheless, we found that the majority of the *Gfap* α and δ mRNA dots were attributed to GFAP processes. The mRNA dots not detected in GFAP intermediate filaments probably belonged to fine distal astrocyte processes devoid of GFAP (e.g. PAPs). These observations suggest that the majority of *Gfap* mRNAs are bound to intermediate filaments, and are consistent with previous reports of colocalization between mRNAs encoding collagens ^31^ and alkaline phosphatase ^32^ on one hand and vimentin (another intermediate filament protein) on the other. Taken as a whole, these findings suggest that intermediate filaments may have crucial roles in the distal distribution of mRNAs. Consequently, it is conceivable that GFAP alterations, deficiency or upregulation (one or the other of which occurs in most neuropathological conditions^17^) might greatly modified the distribution of astrocyte mRNAs and their local translation. In turn, these changes might alter the astrocytes’ functions - particularly at their synaptic and vascular interfaces.

In order to demonstrate the applicability of our approach, we first focused on mRNAs encoding (i) the canonical α isoform of GFAP and (ii) the δ Cter variant, the assembly of which with GFAP α promotes intermediate filament aggregation and dynamic changes ^16, 22^. Interestingly, the results of our experiments in WT mice showed that *Gfap* δ mRNA was more likely than *Gfap* α mRNA to be found in the astrocyte soma. This finding corroborated the results of a previous *in vitro* study in which the proportion of mRNA in primary astrocyte protrusions was higher for *Gfap* α than for *Gfap* δ 33. The high *Gfap* α and δ mRNA density observed in plaque-associated astrocytes was also consistent with previous qPCR-based assays of mRNA in the cortex of APPswe/PS1dE9 mice ^15^. Interestingly, levels of human GFAP α and δ isoforms are elevated in plaque-associated astrocytes in the CA 1-3 region ^34^. Taken together, these results and our observations of elevated mRNA density and distribution in the fine processes of plaque-associated astrocytes suggest that local translation of *Gfap* α and δ mRNA might be a critical mechanism for regulating intermediate filament dynamics in distal astrocyte processes during the progression of AD. Given that the GFAP α/δ isoform ratio is known to strongly influence astrocyte proliferation and malignancy ^35, 36^, our approach might constitute a valuable tool for accurately assessing the differentiation state of astrocytomas in preclinical and clinical settings.

Lastly, we demonstrated that our approach is applicable to any type of mRNA and can also be used in microglia. In fact, the present study is the first to have demonstrated that mRNAs are distributed across microglial processes; this is an important observation in view of the microglia’s complex morphology, motility, and roles in immune surveillance and synaptic remodelling in the brain ^37^. Our results strongly suggest that mRNA distribution and local translation are of physiological significance in this important neural cell type. In conclusion, our new semi-automated *in situ* histological method is the first to have characterized mRNA distribution in astrocytes and microglia.

## Materials and Methods

### Mice

Aldh1L1-Rpl10a mice ^23^ and C57BL6 WT mice were born and then housed under pathogen-free conditions in the animal facility at the Centre Interdisciplinaire de Recherche en Biologie (CIRB, Collège de France, Paris, France). The APPswe/PS1dE9 ^38^ mice were born and housed in the MIRCen animal facility (CEA, Fontenay-aux-Roses, France).

### Ethical approval

All experiments were approved by the French Ministry of Research and Higher Education, and conducted in accordance with the host institution’s ethical standards (Collège de France, Paris, France).

### Aldh1L1:l10a-eGFP TRAP from whole astrocytes and PAPs, and qPCR

Two hippocampi from 5-month-old mice were used for whole-astrocyte polysome extraction. Synaptosomes were prepared from four hippocampi, as described in ^24^ for perisynaptic astrocyte extraction. Polysomes were extracted using the method described in ^39^. Three independent samples were prepared for qPCR analysis. Messenger RNAs were purified using the RNeasy Lipid tissue kit (Qiagen). cDNA was synthesized from 100 ng of whole-astrocyte RNA or the all the PAP mRNA using a Reverse Transcriptase Superscript III kit (Invitrogen) with random primers, and stored at −20°C. Next, 1 µL of cDNA suspension was pre-amplified using SoAmp reagent (BioRad), and droplet qPCR was performed using a QX200™ Droplet Digital™ PCR System (BioRad). The cDNA content was normalized against 45S RNA. TaqMan probes and primer references are listed in Table S1. The data were analyzed by applying a one-way unpaired Mann-Whitney test. The threshold for statistical significance was set to p<0.05.

### Brain slice preparation

Nine-month-old mice were anesthetized with a mix of ketamine/xylazine (0.1 mL/mg) and killed by transcardiac perfusion with 1X phosphate-buffered saline (PBS)/4% paraformaldehyde (PFA) The brain was removed and immersed in 4% PFA overnight at 4°C. The PFA solution was replaced with 15% sucrose for 24 h at 4°C and, lastly, by 30% sucrose for 24 h at 4°C. The brains were cut into 30 µm-thick coronal sections using a Leitz microtome (1400). Sections were stored at −20°C in a cryoprotectant solution (30% glycerol and 30% ethylene glycol in 1X PBS).

### Fluorescent *in situ* hybridization and immunostaining

#### Slice preparation

Slices were carefully washed three times with 1X PBS in a 24-well plate. For the last wash, the 1X PBS was replaced with 7 drops of RNAscope® hydrogen peroxide solution (Advanced Cell Diagnostics Inc.) for 10 min at room temperature (RT); this blocked endogenous peroxidase activity, and resulted in the formation of small bubbles. The slices were washed in Tris-buffered saline with Tween® (50 mM Tris-Cl, pH 7.6; 150 mM NaCl, 0.1% Tween® 20) at RT, and mounted on Super Frost+®-treated glass slides using a paintbrush. Slices were dried at RT for 1 hour in the dark, quickly (in less than 3 s) immersed in deionized water in a glass chamber, dried again for 1 hour at RT in the dark, incubated for 1 h at 60°C in a dry oven, and dried again at RT overnight in the dark.

The slices were rehydrated by rapid immersion (for less than 3 s) in deionized water at RT. Excess liquid was removed with an absorbent paper, and a hydrophobic barrier was drawn. A drop of pure ethanol was applied on the slice for less than 3 s and removed using an absorbing paper. The slides were incubated at 100°C in a steamer, while ensuring that condensation did not fall back on them. A drop of preheated RNAscope® 1X Target Retrieval Reagent (Advanced Cell Diagnostics Inc.) was added to the steamer, and the slides were left for 15 minutes. Next, the slides were washed three times in deionized water at RT, and excess liquid was removed with absorbent paper. A drop of 100% ethanol was applied for 3 minutes, and excess liquid was then removed. A drop of RNAscope® Protease+ solution (Advanced Cell Diagnostics Inc.) was applied and slices were incubated at 40°C in a humid box for 30 minutes. Target retrieval treatment and RNAscope® Protease*+* treatment were used to unmask the mRNAs. Lastly, the slides were washed three times with deionized water at RT.

#### FISH and immunostaining

FISH was performed using the RNAscope® Multiplex Fluorescent Reagent Kit v2 (Advanced Cell Diagnostics Inc.) and specific probes (Table S1), according to the manufacturer’s instructions. Following the FISH procedure, slides were incubated with a blocking solution (5% normal goat serum, 0.375% Triton X-100, and 1 mg.mL^−1^ bovine serum albumin in 1X PBS) for 1 hour at RT, incubated with the primary antibody overnight at 4°C (Table S1), rinsed three times with 1X PBS, and incubated with the secondary antibody (Table S1) for 2h at RT. Lastly, the slides were washed three times in 1X PBS and mounted in Fluor mount and DAPI (Southern Biotech).

### Imaging

Images were acquired using a Yokogawa W1 Spinning Disk confocal microscope (Zeiss) with a 63x oil objective (1.4 numerical aperture). The imaging conditions and acquisition parameters were the same for all slides. The experimental PSF was obtained using carboxylate microsphere beads (diameter: 170 nm; Invitrogen/ThermoFisher Corp.). Except for DAPI, all channels were deconvoluted with Huygens Essential software (version 19.04, Scientific Volume Imaging, The Netherlands; http://svi.nl), using the classic maximum likelihood estimation algorithm and a signal-to-noise ratio of 50 (for the immunofluorescence channel) or 20 (for the FISH channel), a quality change threshold of 0.01, and 150 iterations at most.

### AstroDot and AstroStat

As shown in the Results section, AstroDot can be used to study mRNA density and distribution not only in astrocytes but also in microglia immunolabelled for Iba1. In addition to FISH signals, AstroDot can be used to quantify any type of dot-shaped fluorescence signal. AstroStat was used to analyze the AstroDot results table, using an R script. The programs can be downloaded free of charge from https://github.com/pmailly/Astrocyte_RNA_Analyze and https://github.com/rtortuyaux/astroStat, respectively.

For AstroDot, an image analysis plug-in was developed for the ImageJ/Fiji software ^40, 41^, using Bio-Format (openmicroscopy.org), mcib3D ^42^, GDSC (A. Herbert, https://github.com) and local thickness (B. Dougherty, https://imagej.net/Local_Thickness) libraries.

An ROI enclosing each astrocyte was drawn by hand, using the Fiji polygon tool on the Z projection of the stack. In the ROI Manager, the ROI names were coded as (roi_number-z_top-z_bottom) and saved in a zip file.

#### Plug-in features

The plug-in was designed to process all images in a specific folder containing MetaMorph .nd files, and to read metadata images (channel name, z step, etc.), deconvoluted image channels (except for DAPI), and ROI zip files.

#### AstroDot processing

1. AstroDot’s parameters (the image folder, the channel order, the threshold method, etc.) were displayed in a dialogue box (see Fig. 2C).
2. The immunofluorescence background was estimated using a 0.5 median filter, a binary mask (using Li’s threshold method), and an inversion of the binary mask^43, 44^.The immunofluorescence value was multiplied by the inverted mask and then divided by 255. The background value (bgThreshold) was defined as the mean intensity of all voxels other than those with a value of zero.
3. For each ROI, a substack corresponding to zTop and zBottom (defined in the ROI name) was created for all channels.
4. Semi-automatic determination of the astrocyte or microglial cell nucleus: DAPI fluorescence was processed using a difference of Gaussian filter (kernel: 28–30), a binary mask (using Otsu’s threshold method) and a three-dimensional watershed, to separate nucleus clusters ^45^. An astrocyte nucleus was selected on the basis of its high GFAP immunofluorescence intensity, and was displayed in green. All other detected nuclei were displayed in red. A dialogue box enabled the user to confirm or correct the software’s choice of nuclei.
5. The GFAP immunofluorescence was processed using a 0.5 median filter and a binary mask, using Li’s threshold method. The three-dimensional local thickness of the processes was used to generate a distance map and calculate the local process diameters.
6. FISH dot channel processing: a value of 500 (a manual estimation of the background after deconvolution) was subtracted from each voxel. A difference of Gaussian filter (kernel: 3–1) and a binary mask were applied, using the threshold method defined in the “parameters” dialogue box. The mean dot size volume was computed after the exclusion of dot clusters (volume >2 µm^3^). For dot clusters (which arise when mRNAs are strongly expressed), the dot number was calculated by dividing the cluster by the previously determined mean dot size volume. For each dot, the mean intensity in the Gfap immunofluorescence channel and the distance map value (the process diameter) was calculated.
7. FISH dot classification into three categories
  a. Dot 0 (in red) was a dot in the immunofluorescence background (without using the “Specific mRNA” option only): mean GFAP immunofluorescence intensity ≤ bgThreshold; distance to the boundary of the nucleus > 2 µm.
  b. Dot 1 (in yellow) was a dot in a fine process: mean GFAP immunofluorescence intensity >bgThreshold; distance to the boundary of the nucleus >2 µm; astrocyte process diameter < step in the z calibration (0.3 µm).
  c. Dot 2 (green) was a dot in a large process: mean GFAP immunofluorescence intensity >bgThreshold; distance to the boundary of the nucleus >2 µm; astrocyte process diameter > step in the z calibration (0.3 µm); or a dot in the soma if the distance to the boundary of the nucleus ≤2 µm.
  Hence, Dots 1 and 2 were inside astrocytes, and Dots 0 were outside astrocytes.
8. For each image and for each computed ROI, a .csv output table was generated with the following headers: Image name; ROI name; Background intensity; Astrocyte volume; Dot density inside astrocytes (number of dots 1 + number of dots 2) / astrocyte volume); Percentage of dots outside the astrocyte (number of dots 0 / total dot number); Percentage of dots in astrocyte somata (number of dots less than 2 µm from the boundary of the nucleus / number of dots in astrocytes); Percentage of dots in fine processes (number of dots 1 / number of dots in astrocytes); Percentage of dots in large processes (number of dots 2 – number of dots in somata) / number of astrocyte dots); Mean astrocyte diameter. For each image and each ROI, the selected nucleus, astrocyte channel and classified dot populations were saved as .TIF images.

AstroStat was designed to (i) define the template analysis using a checkbox (working directory, conditions to be compared, paired or unpaired analysis, or data normality plot); (ii) pool data appropriately for each mouse; (iii) test the normality of the data distribution of each group (using Shapiro’s test). If there were more than 30 cells in each group, the central limit theorem was applied; (iv) test the equality of variance (using Fisher’s test) for an unpaired analysis; and (v) compare the means using:

an unpaired analysis:

a. Student’s t-test, for normally distributed data and equal variances
b. Welch-Satterthwaite’s test, for a normal data distribution and unequal variances
c. Wilcoxon’s test for non-normally distributed data

or a paired analysis:

a. Paired Student’s t test for normally distributed data
b. Wilcoxon signed rank test for non-normally distributed data The threshold for statistical significance was set to p<0.05.

## Acknowledgments

This work was funded by a grant from the Fondation pour la Recherche Médicale (AJE20171039094). Noémie Mazaré received a grant from the Fondation pour la Recherche Médicale (FDT201904008077). Romain Tortuyaux received a grant from the “Journées de Neurologie de Langue Francaise”. The creation of the Center for Interdisciplinary Research in Biology (CIRB) was funded by the “Fondation Bettencourt Schueller”. We thank Carole Escartin for the gift of APPswe/PS1dE9 mouse brain slices, and all members of the Orion imaging facility for their high-quality technical support.

## Figure legends

**Supplementary Table S1. Antibodies, qPCR and FISH probes and reagents**

**Supplementary Table S2. Raw data (analyzed using AstroDot and AstroStat) for Gfap α and Gfap δ mRNAs in CA1 and CA3 hippocampal astrocytes from WT and APPswe/PS1dE9 mice**.

SD, standard deviation; N, number of cells analyzed.

