## Supplementary material for "AstroDot: a new method for studying the spatial distribution of mRNA in astrocytes": Supplemantary Table 1

| qPCR probes | | |
|---|---|---|
| **Gene name** | **Supplier** | **Reference** |
| Gfap α RNA (mouse) | BioRad | 10042961 (HEX) dMmuCNS635118061 |
| Gfap δ RNA (mouse) | BioRad | 10042958 (FAM) dMmuCNS795284650 |
| Rpl4 RNA (mouse) | ThermoFisher Scientific | 4331182 (FAM) Mm00834993_g1 |
| 45S RNA (mouse) | ThermoFisher Scientific | 4426961 (FAM) Mm03985792_s1 |

| FISH reagents | | |
|---|---|---|
| **Name** | **Supplier** | **Reference** |
| RNAscope® Multiplex Fluorescent detection reagent V2 | Advanced Cell Diagnostic | 323110 |
| RNAscope® $H_2O_2$ and Protease | Advanced Cell Diagnostic | 322381 |
| Fluoromount-G® | Southern Biotech | 0100-01 |
| Coverglass 0.13 – 0.17mm thick | Immuno Cell | 65.300.13 |
| SuperFrost® Plus slides | VWR | 631-0108 |
| Hydrophobic immunostaining pen | Vector laboratories | H-4000 |

| FISH RNA probes | | | |
|---|---|---|---|
| **Gene name** | **Probe** | **Reference and supplier** | **Dilution** |
| Gfap α RNA | RNAscope® Probe – Mm-Gfap | 313211 Advanced Cell Diagnostic | 1:1[e] |
| Gfap δ RNA | RNAscope® Probe – Mm-Gfap-03 | 557061 Advanced Cell Diagnostic | 1:1[e] |
| Rpl4 RNA | RNAscope® Probe – Mm-Rpl4 | 535821 Advanced Cell Diagnostic | 1:1[e] |

| FISH Fluorophore | | |
|---|---|---|
| **Name** | **Reference and supplier** | **Dilution** |
| Opal 570 | FP14488A Perkin Elmer | 1:1500[e] |
| Opal 650 | FP1496A Perkin Elmer | 1:1500[e] |

| Antibodies | | |
|---|---|---|
| **Name** | **Reference and supplier** | **Dilution** |
| Anti-Glial Fibrillary Acidic Protein antibody produced in rabbit | G9269 Sigma | 1:500[e] |
| Rabbit Anti Iba1 | W1w019-19741 Sobioda | 1:500[e] |
| Goat anti-Rabbit IgG (H+L) Highly Cross-Adsorbed Secondary Antibody, Alexa Fluor 488 | A11034 Invitrogen | 1:1000[e] |
| Goat anti-Rabbit IgG (H+L), Superclonal™ Recombinant Secondary Antibody, Alexa Fluor 647 | A27040 Invitrogen | 1:1000[e] |
