## Supplementary Table 2 for "AstroDot: a new method for studying the spatial distribution of mRNA in astrocytes"

| | | WT | SD | N | Non-Aβ-associated | SD | N | AB-associated | SD | N |
|---|---|---|---|---|---|---|---|---|---|---|
| | | **Gfap α** | | | | | | | | |
| **RNA FISH density** | *CA1* | 1,59 | 0,7 | 175 | 2,11 | 1,21 | 127 | 8,06 | 4,66 | 27 |
| | *CA3* | 1,77 | 0,75 | 94 | 2,25 | 1,3 | 78 | 8,42 | 5,07 | 28 |
| **% RNA in the soma** | *CA1* | 11,49 | 6,77 | 175 | 10,16 | 4,91 | 127 | 7,96 | 4,6 | 27 |
| | *CA3* | 13,27 | 8,15 | 94 | 9,56 | 5,12 | 78 | 7,1 | 3,69 | 28 |
| **% RNA in large processes** | *CA1* | 12,44 | 4,42 | 175 | 11,79 | 4,54 | 127 | 9,61 | 2,88 | 27 |
| | *CA3* | 12,29 | 4,91 | 94 | 10,8 | 4,19 | 78 | 11,6 | 4,09 | 28 |
| **% RNA in fine processes** | *CA1* | 76,07 | 7,56 | 175 | 78,05 | 6,46 | 127 | 82,43 | 5,85 | 27 |
| | *CA3* | 74,45 | 7,61 | 94 | 79,64 | 7,04 | 78 | 81,3 | 5,6 | 28 |
| **% RNA in GFAP+ processes** | *CA1* | 59,48 | 9,04 | 175 | 59,29 | 7,8 | 127 | 60,07 | 7,71 | 27 |
| | *CA3* | 62,16 | 10,05 | 94 | 56,38 | 6,52 | 78 | 59,35 | 8,29 | 28 |
| | | **Gfap δ** | | | | | | | | |
| **RNA FISH density** | | **WT** | **SD** | **N** | **Non-Aβ-associated** | **sd** | **N** | **AB-associated** | **SD** | **N** |
| | | 0,33 | 0,17 | 175 | 0,36 | 0,18 | 127 | 1,16 | 0,76 | 27 |
| **% RNA in the soma** | | 0,35 | 0,17 | 94 | 0,36 | 0,18 | 78 | 1,24 | 0,77 | 28 |
| | | 26,51 | 13,52 | 175 | 26,81 | 11,38 | 127 | 26,98 | 9,66 | 27 |
| **% RNA in large processes** | | 28,5 | 16,39 | 94 | 25,2 | 12,1 | 78 | 24,85 | 14,09 | 28 |
| | | 19,54 | 9,19 | 175 | 17,4 | 7,73 | 127 | 14,08 | 5,73 | 27 |
| **% RNA in fine processes** | | 17,64 | 9 | 94 | 14,79 | 6,84 | 78 | 13,81 | 4,73 | 28 |
| | | 53,95 | 14,67 | 175 | 55,79 | 11,21 | 127 | 58,94 | 9,47 | 27 |
| **% RNA in GFAP+ processes** | | 53,86 | 15,74 | 94 | 60,02 | 10,64 | 78 | 61,34 | 14,5 | 28 |
| | | 74,41 | 11 | 175 | 73,91 | 7,73 | 127 | 75,63 | 7,57 | 27 |
| | | 74,66 | 12,41 | 94 | 69,01 | 8,43 | 78 | 73,66 | 8,75 | 28 |
| | | **WT** | **SD** | **N** | **Non-Aβ-associated** | **SD** | **N** | **AB-associated** | **SD** | **N** |
| **Astrocyte volume (μm$^3$)** | *CA1* | 11541,28 | 4277,51 | 175 | 18264,9 | 6354,5 | 127 | 19550,01 | 8747,74 | 27 |
| | *CA3* | 10430,47 | 3842,5 | 94 | 18181,5 | 5395,3 | 78 | 15144,21 | 6640,57 | 28 |
| **Astrocyte process diameter (mean, in μm)** | *CA1* | 0,32 | 0,03 | 175 | 0,3 | 0,02 | 127 | 0,26 | 0,02 | 27 |
| | *CA3* | 0,32 | 0,04 | 94 | 0,29 | 0,02 | 78 | 0,29 | 0,02 | 28 |
| **Gfap α/ Gfap δ RNA** | *CA1* | 5,22 | 1,7 | 175 | 6,02 | 1,62 | 127 | 7,29 | 1,48 | 27 |
| | *CA3* | 5,25 | 1,42 | 94 | 6,13 | 1,38 | 78 | 7,14 | 2,19 | 28 |

### APPPS1dE9 versus WT

| | | | Non-Aβ-associated / WT | Aβ-associated / WT |
|---|---|---|---|---|
| Astrocyte volume (µm[3]) | CA1 | *Fold Change* | 1,58 | 1,69 |
| | | *p-value* | 1.80E-20 (****) | 3.91E-06 (****) |
| | CA3 | *Fold Change* | 1,74 | 1,45 |
| | | *p-value* | 1.22E-19 (****) | 7.95E-06 (****) |
| Astrocyte process diameter (mean, in µm) | CA1 | *Fold Change* | 0,94 | 0,81 |
| | | *p-value* | 3.16E-13 (****) | 3.23E-08 (****) |
| | CA3 | *Fold Change* | 0,91 | 0,91 |
| | | *p-value* | 3.21E-08 (****) | 1.04E-04 (***) |
| Gfap α/ Gfap δ RNA | CA1 | *Fold Change* | 1,15 | 1,40 |
| | | *p-value* | 5.05E-05 (****) | 2.05E-08 (****) |
| | CA3 | *Fold Change* | 1,17 | 1,36 |
| | | *p-value* | 6.12E-05 (****) | 1.34E-04 (***) |

| | | | Gfap α | | Gfap δ | |
|---|---|---|---|---|---|---|
| | | | Non-Aβ-associated / WT | Aβ-associated / WT | Non-Aβ-associated/ WT | Aβ-associated / WT |
| RNA FISH density | CA1 | *Fold Change* | 1,33 | 5,07 | 1,09 | 3,52 |
| | | *p-value* | 2.27E-05 (****) | 3.66E-16 (****) | 1.50E-01 (NS) | 8.00E-14 (****) |
| | CA3 | *Fold Change* | 1,27 | 4,76 | 1,03 | 3,54 |
| | | *p-value* | 4.81E-03 (**) | 3.77E-12 (****) | 6.85E-01 (NS) | 7.94E-10 (****) |
| % RNA in the soma | CA1 | *Fold Change* | 0,88 | 0,69 | 1,01 | 1,02 |
| | | *p-value* | 4.93E-02 (*) | 3.86E-03 (**) | 8.35E-01 (NS) | 8.27E-01 (NS) |
| | CA3 | *Fold Change* | 0,72 | 0,54 | 0,88 | 0,87 |
| | | *p-value* | 3.67E-04 (***) | 5.53E-05 (****) | 1.31E-01 (NS) | 3.99E-01 (NS) |
| % RNA in large processes | CA1 | *Fold Change* | 0,95 | 0,77 | 0,89 | 0,72 |
| | | *p-value* | 2.10E-01 (NS) | 9.07E-04 (***) | 2.92E-02 (*) | 1.34E-03 (**) |
| | CA3 | *Fold Change* | 0,88 | 0,94 | 0,84 | 0,78 |
| | | *p-value* | 3.53E-02 (*) | 4.42E-01 (NS) | 1.95E-02 (*) | 1.71E-02 (*) |
| % RNA in fine processes | CA1 | *Fold Change* | 1,03 | 1,08 | 1,03 | 1,09 |
| | | *p-value* | 1.76E-02 (*) | 1.27E-05 (****) | 2.18E-01 (NS) | 2.83E-02 (*) |
| | CA3 | *Fold Change* | 1,07 | 1,09 | 1,11 | 1,14 |
| | | *p-value* | 7.53E-06 (****) | 1.24E-05 (****) | 2.73E-03 (**) | 2.67E-02 (*) |
| % RNA in GFAP+ processes | CA1 | *Fold Change* | 1,00 | 1,01 | 0,99 | 1,02 |
| | | *p-value* | 8.54E-01 (NS) | 7.46E-01 (NS) | 6.42E-01 (NS) | 5.16E-01 (NS) |
| | CA3 | *Fold Change* | 0,91 | 0,95 | 0,92 | 0,99 |
| | | *p-value* | 1.02E-05 (****) | 1.80E-01 (NS) | 5.30E-04 (***) | 4.30E-01 (NS) |

### CA1 versus CA3

| | | WT | Non-Aβ-associated | AB-associated |
|---|---|---|---|---|
| **Astrocyte volume (µm$^3$)** | ***Fold Change*** | 1,11 | 1,00 | 1,29 |
| | ***p-value*** | 3.57E-02 (*) | 9.23E-01 (NS) | 6.01E-02 (NS) |
| **Astrocyte process diameter (mean, in µm)** | ***Fold Change*** | 1,00 | 1,03 | 0,90 |
| | ***p-value*** | 3.91E-01 (NS) | 2.61E-02 (*) | 5.87E-01 (NS) |
| **Gfap α/ Gfap δ RNA** | ***Fold Change*** | 0,99 | 0,98 | 1,02 |
| | ***p-value*** | 9.08E-01 (NS) | 6.27E-01 (NS) | 7.67E-01 (NS) |

| | | **Gfap α** | | | **Gfap δ** | | |
|---|---|---|---|---|---|---|---|
| | | **WT** | **Non-Aβ-associated** | **AB-associated** | **WT** | **AB-not associated** | **AB-associated** |
| **RNA FISH density** | ***Fold Change*** | 0,90 | 0,94 | 0,96 | 0,94 | 1,00 | 0,94 |
| | ***p-value*** | 5.26E-02 (NS) | 4.55E-01 (NS) | 8.61E-01 (NS) | 2.21E-01 (NS) | 7.49E-01 (NS) | 6.46E-01 (NS) |
| **% RNA in the soma** | ***Fold Change*** | 0,87 | 1,06 | 1,12 | 0,93 | 1,06 | 1,09 |
| | ***p-value*** | 7.15E-02 (NS) | 4.05E-01 (NS) | 5.99E-01 (NS) | 3.16E-01 (NS) | 3.37E-01 (NS) | 5.18E-01 (NS) |
| **% RNA in large processes** | ***Fold Change*** | 1,01 | 1,09 | 0,83 | 1,11 | 1,18 | 1,02 |
| | ***p-value*** | 7.95E-01 (NS) | 1.21E-01 (NS) | 1.12E-01 (NS) | 1.04E-01 (NS) | 1.50E-02 (*) | 1.00E00 (NS) |
| **% RNA in fine processes** | ***Fold Change*** | 1,02 | 0,98 | 1,01 | 1,00 | 0,93 | 0,96 |
| | ***p-value*** | 9.36E-02 (NS) | 1.01E-01 (NS) | 4.67E-01 (NS) | 9.65E-01 (NS) | 8.13E-03 (**) | 4.70E-01 (NS) |
| **% RNA in GFAP+ processes** | ***Fold Change*** | 0,96 | 1,05 | 1,01 | 1,00 | 1,07 | 1,03 |
| | ***p-value*** | 2.60E-02 (*) | 6.36E-03 (**) | 7.39E-01 (NS) | 8.67E-01 (NS) | 3.20E-05 (****) | 2.89E-01 (NS) |
